# A Microfluidic Transistor for Liquid Signal Processing

**DOI:** 10.1101/2023.05.31.543146

**Authors:** Kaustav A. Gopinathan, Avanish Mishra, Baris R. Mutlu, Jon F. Edd, Mehmet Toner

**Affiliations:** BioMEMS Resource Center, Center for Engineering in Medicine and Surgical Services, Massachusetts General Hospital, Boston, MA 02114, USA; Cancer Center, Massachusetts General Hospital, Boston, MA 02114, USA; Department of Surgery, Massachusetts General Hospital, Boston, MA 02129, USA; Shriners Hospitals for Children, Boston, MA 02114, USA

## Abstract

Microfluidics have enabled significant advances in molecular biology^1–3^, synthetic chemistry^4,5^, diagnostics^6,7^, and tissue engineering^8^. However, there has long been a critical need in the field to manipulate fluids and suspended matter with the precision, modularity, and scalability of electronic circuits^9–11^. Just as the electronic transistor enabled unprecedented advances in the control of electricity on an electronic chip, a microfluidic analogue to the transistor could enable improvements in the complex, scalable control of reagents, droplets, and single cells on an autonomous microfluidic chip. Prior works on creating a microfluidic analogue to the electronic transistor^12–14^ could not replicate the transistor’s saturation behavior, which is crucial to perform analog amplification^15^ and is fundamental to modern circuit design^16^. Here we exploit the fluidic phenomenon of *flow-limitation*^17^ to develop a microfluidic element with flow-pressure characteristics completely analogous to the current-voltage characteristics of the electronic transistor. As this microfluidic transistor successfully replicates all of the key operating regimes of the electronic transistor (linear, cut-off and saturation), we are able to directly translate a variety of fundamental electronic circuit designs into the fluidic domain, including the amplifier, regulator, level shifter, logic gate, and latch. Finally, we demonstrate a “smart” particle dispenser that senses single suspended particles, performs liquid signal processing, and accordingly controls the movement of said particles in a purely fluidic system without electronics. By leveraging the vast repertoire of electronic circuit design, microfluidic transistor-based circuits are easy to integrate at scale, eliminate the need for external flow control, and enable uniquely complex liquid signal processing and single-particle manipulation for the next generation of chemical, biological, and clinical platforms.

---

As the precision and complexity requirements of microfluidic applications grow, controlling the flow of reagents, droplets, and cells on these platforms has become increasingly challenging^9–11^. Microfluidic platforms have conventionally addressed the problem of complex flow control with embedded electromechanical or pneumatic valves actuated by external computer controllers^18–20^. However, these external control systems are challenging to scale and limit the complexity of the fluidic operations that can be performed^1,9,10^. More recent work has resulted in the development of miniature self-actuated valves that function without an external controller^12,13^. While the valves in these systems are capable of switching flows on or off in simple fluidic circuits, they do not replicate the saturation behavior of the transistor, and cannot amplify an analog signal—the defining feature of the transistor^15,21^. Without this characteristic saturation behavior, these microfluidic valves cannot be used to perform analog signal processing, and limit the potential of applying modular, complex circuit designs from electronics towards the scalable control of liquids on a chip.

The key technical advance of this study is to exploit the fluidic phenomenon of *flow-limitation* to create a microfluidic transistor that replicates all of the electronic transistor operating regimes (linear, cut-off, and saturation). After characterizing this microfluidic transistor, we use it to demonstrate microfluidic analogues for several classic electronic building blocks including the amplifier, regulator, level shifter, NAND gate, and SR-latch. These circuit blocks enable modular liquid signal processing on-chip without external controllers. Finally, to showcase the microfluidic transistor’s ability to directly manipulate particles suspended in fluids, we demonstrate a “smart” particle dispenser. This dispenser is capable of sensing single suspended particles, performing signal processing operations, and dispensing the particles in a controllable manner. We integrate this dispenser along with several other microfluidic transistor-based circuit blocks into a self-contained system that automatically performs deterministic single-particle ordering and concentration without any external optical or electronic components.

The microfluidic transistor consists of two crossed channels of fluid separated by a deformable membrane (Fig. 1a) and is fabricated entirely from elastomer using standard soft-lithography techniques (see Methods). It is represented schematically in Fig. 1b, where the flow and pressures relevant to its function are also labelled. When a pressure difference *P_SD_* is applied between the source and drain terminals, the membrane between the crossed channels deforms itself. With carefully chosen channel geometry, this self-deformation limits the volumetric flow *Q* passing through the drain in a special non-linear manner known as flow-limitation. The extent of this effect and thus the flow rate through the device can be modulated by applying a pressure *P_GS_* between the gate and source terminals.

**Fig. 1.**
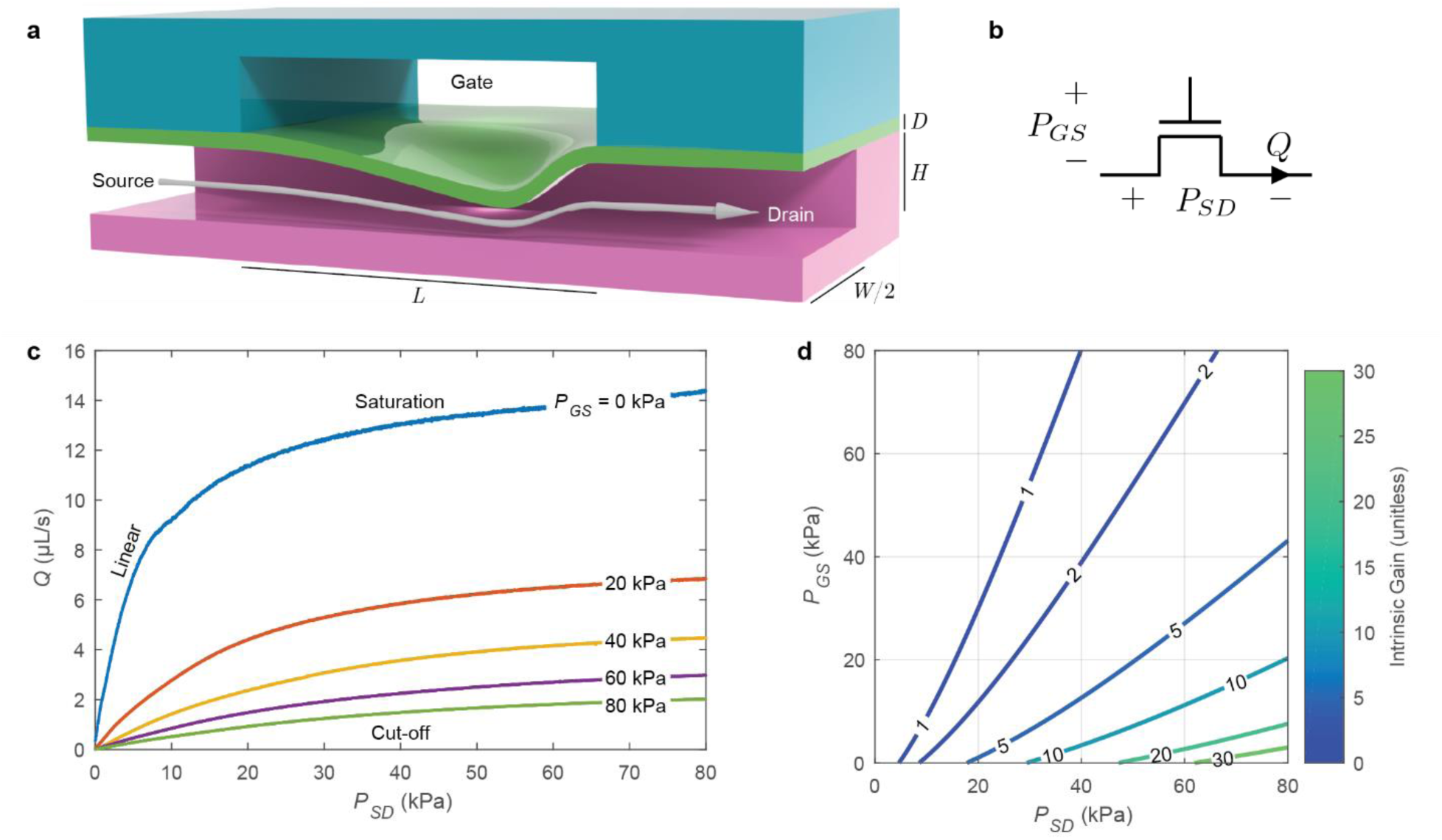
Elastomeric channels exhibit pressure-controlled flow-limitation analogous to an electronic transistor. **a,** Longitudinal section of the microfluidic transistor, fabricated from two layers of thick elastomer with channels (magenta, teal) and a thin elastomer membrane (green). Pressure applied between the gate and the source deflects the membrane, restricting flow (arrow) from the source to the drain in a non-linear fashion known as flow-limitation. **b,** Schematic symbol for the microfluidic transistor. The pressure difference between the gate and the source is *P_GS_*, and the pressure difference between the source and the drain is *P_SD_*. Volumetric flow through the drain is *Q*. **c,** Experimentally measured characteristic curves of the microfluidic transistor, demonstrating all three operation regimes seen in electronic transistors (linear, cut-off, and saturation). **d,** Contour plot of the intrinsic gain of the microfluidic transistor as a function of *P_GS_* and *P_SD_*, depicting a large region with intrinsic gain greater than one.

The microfluidic transistor is characterized in a fashion analogous to that of the electronic field-effect transistor. Figure 1c provides the characteristic curves for a microfluidic transistor with dimensions provided in Extended Data Table 1. Volumetric flow *Q* is recorded while *P_SD_* is swept across a range of pressures and *P_GS_* is held at fixed values, resulting in the fluidic version of the classic transistor characteristic curves. For visual clarity, a subset of the measured characteristic curves is plotted in Fig. 1c, and the complete set of curves for all measured values of *P_GS_* is provided in Extended Data Fig. 1a. Additional transconductance characterization of the microfluidic transistor is provided in Extended Data Fig. 1b.

A defining characteristic for any transistor is its *intrinsic gain*, a dimensionless measure of the maximum analog amplification achievable for a given set of potentials applied across the source, gate, and drain^15^. Crucially, for a microfluidic element to amplify like a transistor, there must be a practically achievable range of values for *P_SD_* and *P_GS_* where the intrinsic gain is greater than one (Supplementary Text I). Figure 1d shows a contour plot of the intrinsic gain as a function of applied *P_GS_* and *P_SD_*, computed using the characterization data of Extended Data Fig. 1a. The contour plot reveals a large operating region where the intrinsic gain is much greater than one, indicating that this microfluidic element exhibits true transistor behavior and is capable of greatly amplifying analog signals at a wide range of applied operating pressures.

The key to achieving high intrinsic gains in these microfluidic transistors is to exploit the phenomenon of flow-limitation. This phenomenon is observed in certain confined flows through tubes with deformable boundaries (including the human *vena cava*), where increasing the pressure drop across the tube beyond a threshold does not substantially increase the flow rate through the tube^17,22^. This flow-limitation phenomenon occurs in systems where the dimensionless Shapiro number is greater than one^23^. For the microfluidic channels considered here, the Shapiro number *S* is given by (Supplementary Text II):

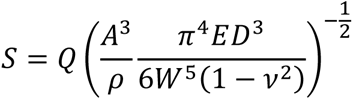

Where *Q* is the flow rate, *ρ* is the fluid density, *A* is the channel cross-sectional area, *W* is the channel width, *D* is the membrane thickness, *E* is the membrane Young’s modulus, and *v* is the membrane Poisson ratio. Using the above equation (valid when *P_GS_* = 0) and the measured data from Fig. 1c, we observe that when the Shapiro number exceeds one, the flow-pressure characteristic of the transistor diverges from the typical linear Poiseuille behavior and enters flow-limitation (Extended Data Fig. 2). This may also be observed in the “Saturation” region of Fig 1c where the flow-pressure curves become flat. The flow-limitation effect observed in the microfluidic transistor is similar in practice to the saturation behavior of the field-effect transistor, and these effects are the fundamental mechanisms by which the transistor devices achieve their high intrinsic-gains and perform amplification.

**Fig. 2.**
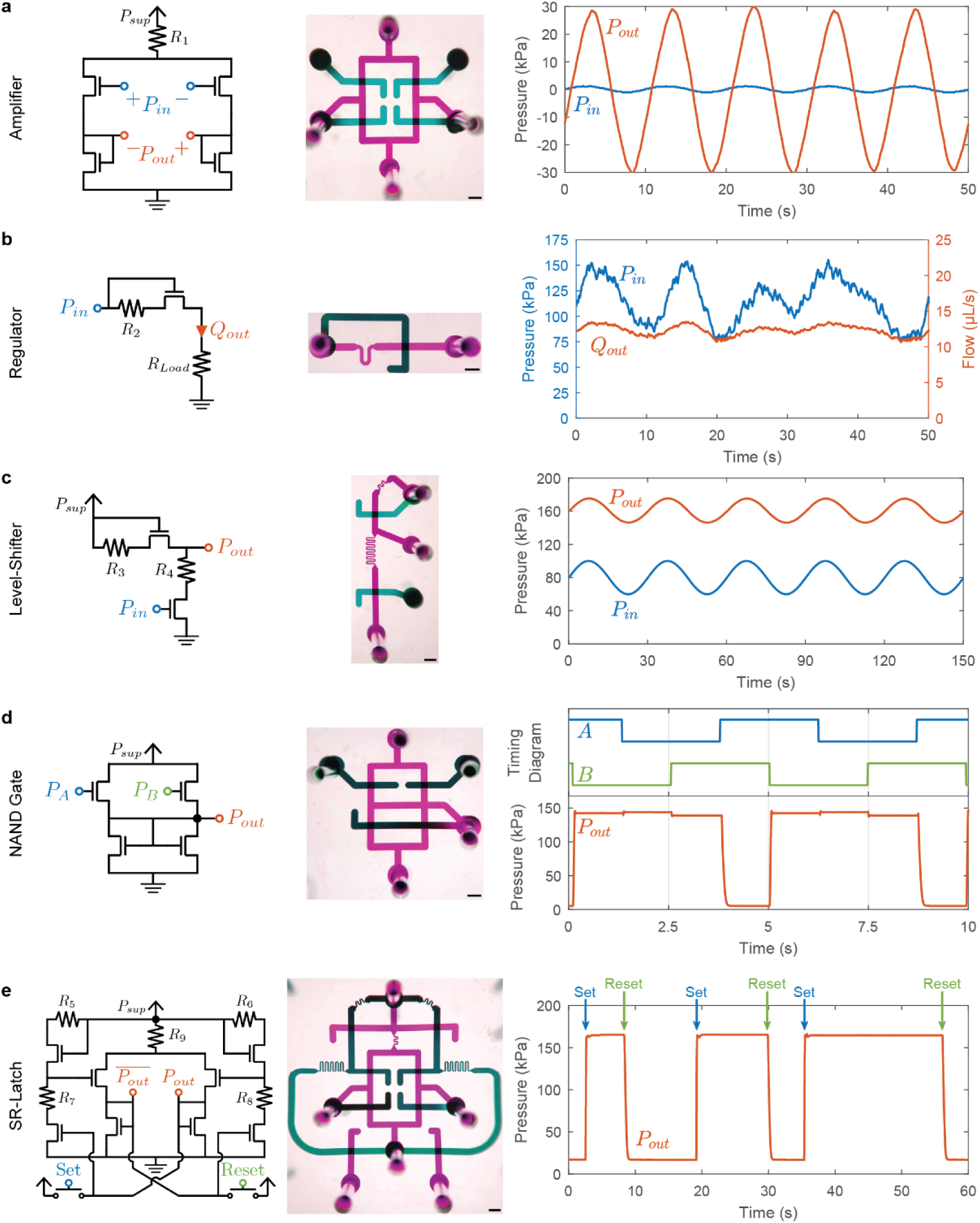
Microfluidic transistor-based circuits replicate the behavior of classic electronic circuits. For each circuit, the schematic diagram (left), a photo of the microfluidic implementation (middle, false color, scale bars 1 mm), and a representative demonstration of circuit function (right) is provided. **a,** A fluidic amplifier. The input pressure signal (blue) is amplified with a gain of 22 to generate the output signal (orange). **b,** A flow regulator. The input pressure (blue) varying from 75 to 150 kPa is regulated to supply a target flow (orange) of 12±1.5 μL/s to a load. **c,** A level shifter. The baseline of a varying input pressure signal (blue) is shifted up by 80 kPa to produce an output pressure signal with the same morphology (orange). **d,** A NAND gate. The output signal (orange) is low only if both input signals (blue and green) are high. **e,** An SR-latch. The persistent state of the latch (orange) can be set to high or low pressure based on transient pulses applied to Set (blue) or Reset (green).

To illustrate the flexibility of the microfluidic transistor and the ease with which it can be directly substituted into pre-existing designs from electronics, we demonstrate the microfluidic analogues to a range of classic electronic circuit blocks (Fig. 2). For each of these circuits, we also provide several characterization studies to evaluate their performance, similar to the studies typically found in electronic datasheets (Extended Data Fig. 3-5). The specific component values used in these circuits are provided in Extended Data Table 1.

**Fig. 3.**
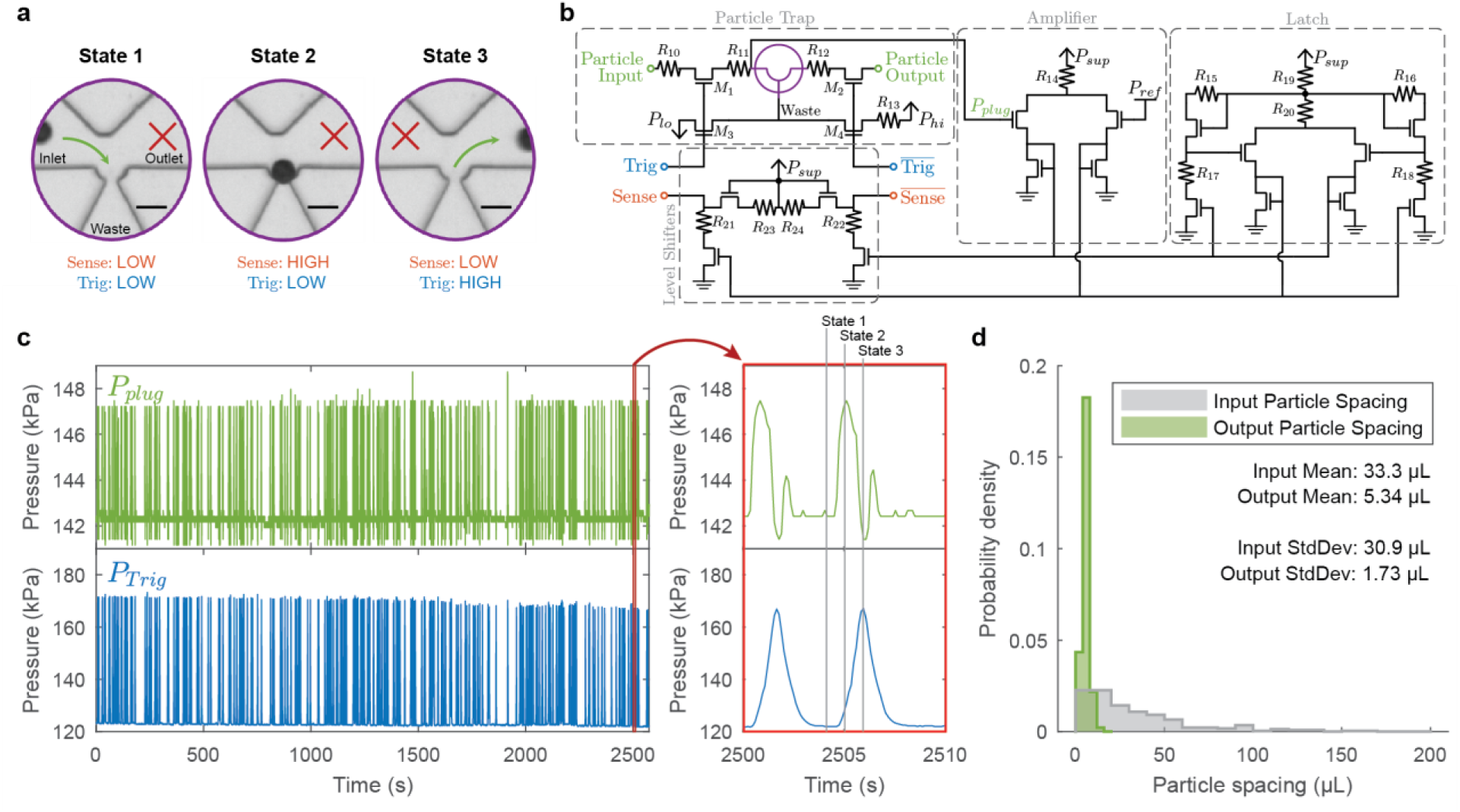
Microfluidic transistors enable smart dispenser circuits for autonomous single particle manipulation. **a,** Overview of the smart dispenser operation, depicting the core microfluidic trap in different states as it senses and dispenses a single particle (scale bars 50 μm). **b,** Circuit schematic of the smart dispenser comprising of several circuit blocks and the microfluidic trap (purple). **c,** Deterministic single-particle ordering and concentration using the smart dispenser. This dispenser configuration has the Trig and Sense lines directly connected, so that individual particles are sensed and immediately dispensed into the output channel. Pressure signals from the trap itself (*P_plug_*) and the trigger (*P_Trig_*) for a run of n=230 particles are shown, along with a representative dispense event to observe the individual dynamics (red inset). **d,** Histograms of input and output particle spacing when using the smart dispenser in this configuration, showing a 6-fold drop in the spacing mean (indicating particle concentration) and a 17-fold drop in the spacing standard deviation (indicating particle ordering).

Since amplification is the defining characteristic of a transistor^21^, we first demonstrate microfluidic transistors in a differential amplifier (Fig. 2a). This analog circuit is capable of amplifying an input differential pressure signal by a gain factor of over 20. Additional characterization of the frequency response, common-mode rejection, and distortion for this circuit are provided in Extended Data Fig. 3. Amplifiers are the fundamental building blocks of analog circuits, used ubiquitously in signal processing and feedback control^15,16^. They are also used as buffers in digital logic.

A flow regulator is demonstrated in Fig. 2b. This analog circuit is designed to supply a constant output flow to a downstream load regardless of the input pressure level. Additional characterization of the load and line regulation for this circuit are provided in Extended Data Fig. 4a-b. Regulators may be used to stabilize flow when microfluidic devices are supplied by unregulated pressure sources (such as balloons or hand pumps) in mobile settings.

A level shifter is demonstrated in Fig. 2c. This analog circuit is designed to translate the baseline pressure of the input signal to a higher output baseline pressure without affecting the morphology of the signal. Additional characterization of the shift amount and gain for this circuit are provided in Extended Data Fig. 4c-d. Level shifters allow multiple circuit blocks to be cascaded after each other, even if they require different biasing pressures, enabling design modularity.

A NAND gate is demonstrated in Fig. 2d. This digital logic gate produces a low output pressure only if both inputs are at a high pressure. NAND gates are universal logic gates, so can be combined to implement all other boolean logic operations (AND, OR, XOR, etc.) for general digital signal processing. Additional characterization of the output dynamics and transfer characteristics for this circuit are provided in Extended Data Fig. 5a-d. Logic gates may be used to synchronize fluidic events, perform sequential fluidic operations, or even compute binary arithmetic.

An SR-latch (bistable multivibrator) is demonstrated in Fig. 2e. This digital circuit has two stable output states that can be set high or low persistently after receiving a transient “set” or “reset” pressure pulse, and therefore hold memory. This circuit is composed of a cross-coupled differential amplifier with two level shifters, and so also illustrates how the previously described building blocks can be combined to perform more complex operations. Additional characterization of the set and reset response dynamics for this circuit is provided in Extended Data Fig. 5e-f. Cascaded latches act as fluidic memory and are capable of storing binary numbers. As a result, they may be used to count fluidic events or perform sequential combinatorial operations that require memory of the circuit’s previous state.

While the circuits of Fig. 2 demonstrate how the microfluidic transistor can be used to replicate the major building blocks of electronics, we sought to also demonstrate a unique application for the microfluidic transistor that cannot be performed by an electronic transistor: directly manipulating single particles suspended in liquid. Figure 3 demonstrates a “smart” particle dispenser capable of detecting and programmatically dispensing individual suspended particles. At the core of the dispenser is a microfluidic particle trap with an inlet, outlet, and waste channel (Fig. 3a). Normally, when there is no particle in the trap, fluid flows directly from the inlet to the waste channels (State 1). When a particle becomes trapped, the dispenser detects its presence and produces a high Sense pressure signal, indicating that it is holding a trapped particle and is awaiting the trigger signal to dispense it (State 2). If the dispenser then receives a high Trig pressure signal, the trapped particle is ejected into the outlet channel (State 3). After the particle is dispensed, the dispenser returns to its initial state to accept a new particle.

To perform this complex sequence of dispensing operations, several signal processing circuit blocks from Fig 2 are utilized in the dispenser’s control circuitry (Fig 3b). When a particle is trapped, the pressure upstream of the trap (*P_plug_*) rises slightly. An amplifier circuit block is used to amplify this small change and compare it with a reference threshold pressure, producing a pair of complementary signals indicating the presence of a particle. The latch circuit block ensures complementarity of the signals and suppresses any spurious noise events that were amplified. Finally, these signals are shifted up using level shifter circuit blocks to produce the output Sense and complementary 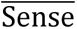 signals. The complementary Trig and 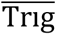 signals are used to control the direction of flow in the trap. Details on the specific component values, sizes, and pressures used here are provided in Extended Data Table 1.

This smart dispenser offers numerous applications for counting, ordering, encapsulating, and distributing individual particles or biological cells. Here we demonstrate a simple configuration of the dispenser circuit block by connecting the Sense and Trig lines directly to each other. This configuration results in deterministic particle ordering and concentration in the output channel, as demonstrated in Fig. 3c using 40 μm polystyrene beads. While particles at the dispenser input are randomly longitudinally spaced (as a Poisson process)^24^, particles in the dispenser output channel follow a tight distribution of equal spacing along the output stream (Fig. 3d, Extended Data Fig. 6). The 6-fold drop in the spacing mean and the 17-fold drop in the spacing standard deviation indicate that this configuration of the smart dispenser successfully concentrated and ordered the particles. It is important to note that all the liquid signal processing, particle manipulation, and flow control demonstrated here were performed entirely on-chip through judicious use of amplifiers and other microfluidic transistor-based circuit blocks, requiring only constant pressure sources to supply power.

Although here we illustrated a simple configuration of the dispenser for particle ordering, additional microfluidic transistor-based circuitry (such as those described in Fig. 2) may be readily integrated for more advanced logic on the Sense and Trig signals to accomplish more complex particle dispensing tasks such as synchronization between multiple dispensers for multi-particle encapsulation or combinatorial operations.

Using microfluidic transistors for complex flow control has several significant advantages over prior methodologies. Many microfluidic systems use external computers that switch on or off individual electromechanical or pneumatic valves embedded in the microfluidic chip^18–20^. Unfortunately, these off-chip approaches are practically limited in complexity and scalability due to the difficulty in interfacing the microfluidic chip with the external electronics and the large footprint of the many external electrical or pneumatic control lines needed^9,10^. Performing signal processing entirely in the fluidic domain circumnavigates this key deficiency.

Our approach to complex flow control is also fundamentally distinct from the existing repertoire of integrated valve-based microfluidic systems. Building off pioneering advances from the Quake^18,25^ and Mathies^14^ groups, recent valve-based systems use movable membranes (“switch valves”)^9,12^ or embedded rigid disks (“gain valves”)^13,26^ to control the flow of fluid through a channel. Unlike the microfluidic transistors described in this work, microfluidic valves do not exploit the flow-limitation phenomenon and instead rely on physically blocking the flow channel by moving a sealing surface. As a result, these valves behave akin to an electronic relay that switches flows entirely on or off. These valves have previously been used to make a number of digital circuits, including logic gates, clocks, and latches^9,13^. However, since these valves do not replicate the saturation regime of the transistor and do not have an intrinsic gain, they cannot be used to build a large class of analog circuits for signal processing, as they cannot replicate the amplifier, regulator, level shifter, and smart particle dispenser shown here. Additionally, these sealing-surface valves suffer from unequal opening and closing pressure thresholds that can even vary based on surrounding circuitry, making circuit design challenging and limiting the scale of the digital circuit designs they could be used for as well^27^.

The microfluidic transistor described here replicates all the operational regimes of the electronic transistor and demonstrates a large region of high intrinsic gain suitable for both analog signal processing and digital logic in one device. It can be easily fabricated and used to directly translate a number of classic building block circuits from analog and digital electronics, functioning without any external control pneumatics, electronics, or optics. Thanks to this self-contained nature, these modular circuit blocks may be readily scaled using amplifiers and level shifters to perform manipulations on liquids and particles with unprecedented complexity. With the ability to both process liquid signals and manipulate single particles based on those signals, we envision that microfluidic transistor-based circuitry will unlock the breadth and depth of electronic circuit design for the next generation of biological and chemical processing.

## Methods

### Microfluidic device fabrication

All devices used in this work were fabricated from two layers of polydimethylsiloxane (PDMS) and a thin silicone membrane (Fig. 1a). Standard soft-lithography techniques were used to fabricate each layer. In brief, SU-8 50 negative photoresist (Kayaku Advanced Materials, Inc.) was spin-coated onto a silicon wafer at 2450 rpm for 30 sec. The channels were patterned onto the SU-8 by exposing the wafer with 365 nm UV radiation through a photomask. The wafer was subsequently developed using Baker BTS-220 SU-8 developer to create the mold for the PDMS. For each device, two such molds were made for the upper and lower PDMS layers. PDMS (Dow Sylgard 184 Kit, Ellsworth Adhesives) was prepared in a 6:1 ratio of base to cross-linker and poured into each mold to create a 4 mm thick layer. The high ratio of cross-linker to base was used to minimize the deformation of the PDMS resistor channels as the channels were pressurized. The PDMS layers were cured in a convection oven for 20 hours at 70°C, then cut and peeled from the mold.

After casting the upper and lower layers of the device from PDMS, they were assembled to make the final microfluidic chips (Extended Data Fig. 7). A 1.2 mm biopsy punch was used to punch out specific ports in the upper PDMS layer. The PDMS layer was then bonded to a 20 μm thick silicone membrane (Elastosil Film 2030 250/20, Wacker Chemie AG) via oxygen plasma treatment and baked at 80°C for 15 minutes on a hotplate. A 1.2 mm biopsy punch was then used to create the remaining ports in the bonded assembly of the upper layer and membrane. The membrane side of the assembly was then bonded to the lower PDMS layer via oxygen plasma treatment and baked at 90°C for 15 minutes on a hotplate. The higher temperature ensured sufficient heat reached the bonding surfaces through the lower PDMS layer.

### Device setup and testing

All devices were primed by submerging the device under distilled water and applying a vacuum of approximately 75 kPa below atmosphere for 10 minutes. Air was then slowly released into the vacuum chamber while the devices were submerged, priming the channels with distilled water. Unless otherwise specified, all fluidic connections were made with 0.03-inch inner diameter fluorinated ethylene propylene (FEP) tubing (1520XL, IDEX-HS) and PEEK fittings purchased from IDEX Health & Sciences. The various tubular fluidic resistors were made using 0.01-inch inner diameter FEP tubing (1527L, IDEX-HS). The specific resistor lengths and other component details for each circuit are provided in Extended Data Table 1. Computer-controlled pressure sources (LineUp FlowEZ, Fluigent) were used to supply pressures for characterization of the microfluidic devices. Unless otherwise specified, all reservoirs for the pressure sources (P-CAP, Fluigent) were filled with 1X phosphate-buffered saline (PBS) (Gibco PBS, Fisher Scientific). All pressure measurements were made using Honeywell pressure sensors (ABPDRRV015PDAA5) and logged on a computer using MATLAB. All flow measurements were made using Sensirion flow meters (SLI-1000).

### Single transistor characterization

The pinout for the single transistor chip is given in Extended Data Figure 8a. Extended Data Figure 8b provides the setup used to measure the transistor characteristic curves (Fig. 1c, Extended Data Fig. 1a). The “Gate” pressure source and the “Channel” pressure source used a Fluigent LU-FEZ-2000 module and a Fluigent LU-FEZ-1000 module respectively to control the pressure. To apply a given *P_SD_* and *P_GS_* to the device, the pressure at “Channel” was set to *P_SD_* and the pressure at “Gate” was set to *P_GS_* + *P_SD_*. To generate the characteristic curves, *P_GS_* was set to 0 kPa, *P_SD_* was swept from 0 kPa to 80 kPa over the course of 600 s, and the flow *Q* was recorded to generate each curve. Then, *P_GS_* was incremented by 5 kPa and the process was repeated until *P_GS_* reached 80 kPa.

To obtain the intrinsic gain contour plot (Fig. 1d), the two-dimensional surface of points collected from the previous characteristic curve measurements was smoothed using a two-variable rational polynomial function of degree one in the numerator and degree two in the denominator. The smoothed polynomial was confirmed to fit the raw data well (*R*^2^ > 0.99) and was used to avoid noise when computing the numerical derivatives. The intrinsic gain was then calculated in MATLAB from the smoothed data (Equation 1.3, Supplementary Information I).

The same setup (Extended Data Fig. 8b) was used to measure the transistor transfer characteristics (Extended Data Fig. 1b). To generate the transfer characteristic curves, *P_SD_* was set to 20 kPa, *P_GS_* was swept from 0 kPa to 80 kPa over the course of 300 s, and the flow *Q* was recorded to generate each curve. Then, *P_SD_* was incremented by 20 kPa and the process was repeated until *P_SD_* reached 80 kPa.

### Amplifier characterization

The pinout for the amplifier is given in Extended Data Figure 8c. Extended Data Figure 8d provides the setup used to demonstrate the amplifier (Fig. 2a). The “Supply” pressure source used a Fluigent LU-FEZ-7000 module to control the pressure. The “Input1” and “Input2” pressure sources used two Fluigent LU-FEZ-2000 modules. The tubing dimensions used for the resistances are provided in Extended Data Table 1. The “Supply” pressure source was set to 250 kPa. The “Input1” and “Input2” pressure sources applied a common-mode bias of 175 kPa and a differential sinusoidal signal of amplitude 1 kPa and a period of 10 s. The differential input and output signals were measured by pressure sensors.

The same setup (Extended Data Fig. 8d) was used to measure the amplifier distortion (Extended Data Fig. 3a). The “Supply” pressure source was set to 250 kPa. Over the course of 150 s, the “Input1” pressure source was swept from 180 kPa to 170 kPa and the “Input2” pressure source was swept from 170 kPa to 180 kPa. The differential input and output signals were measured by pressure sensors.

Extended Data Figure 8e provides the setup used to measure the amplifier common-mode rejection (Extended Data Fig. 3b). The “Supply” and “Input” pressure sources used a Fluigent LU-FEZ-7000 and a Fluigent LU-FEZ-2000 module respectively to control the pressure. The tail resistance (*R*_1_) was made using 30 cm of 0.01 in diameter FEP tubing (1527L, IDEX-HS). The “Supply” pressure source was set to 250 kPa and the “Input” pressure source was swept from 160 kPa to 200 kPa over the course of 150 s. The differential output signal was measured by a pressure sensor.

Extended Data Figure 8f provides the setup used to determine the amplifier frequency response (Extended Data Fig. 3c). The “Supply” pressure source used a Fluigent LU-FEZ-7000 module to control the pressure. The “InHigh” and “InLow” pressure sources used two Fluigent LU-FEZ-2000 modules. The “Switch” was a Fluigent 2-switch (2SW002). The tail resistance (*R*_1_) was made using 30 cm of 0.01 in diameter FEP tubing (1527L, IDEX-HS). The “Supply” pressure source was set to 250 kPa, the “InLow” pressure source was set to 175 kPa, and the “InHigh” pressure source was set to 177 kPa. The “Switch” was set to toggle every 15 s. The differential input and output signals were measured by pressure sensors and data was collected over 500 s.

To generate the frequency response plot of the amplifier (Extended Data Fig. 3c), the differential input and output signals were resampled to a constant sampling frequency, then converted to the frequency domain. Since a square wave excitation signal in the time domain only produces odd harmonics in the frequency domain, the first 40 odd harmonics of the input and output frequency-domain signals were used to generate the frequency response plot points.

### Flow regulator characterization

The pinout for the regulator chip is given in Extended Data Figure 8g. Extended Data Figure 8h provides the setup used to demonstrate the flow regulator (Fig. 2b). The “Input” pressure source used a Fluigent LU-FEZ-2000 module to control the pressure. The *R_load_* resistance was made using 20 cm of 0.01 in diameter FEP tubing (1527L, IDEX-HS). To simulate a poorly regulated pressure source, the “Input” pressure source was applied an arbitrary randomly-generated pressure waveform ranging from approximately 75 kPa to 150 kPa over the course of 50 s while the flow through the load was recorded.

The same setup (Extended Data Fig. 8h) was used to measure the line regulation of the flow regulator (Extended Data Fig. 4a). The *R_load_* resistance was made using 20 cm of 0.01 in diameter FEP tubing (1527L, IDEX-HS). The “Input” pressure source was swept from 0 kPa to 150 kPa over the course of 300 s and the flow was recorded.

Extended Data Figure 8i provides the setup used to measure the load regulation of the flow regulator (Extended Data Fig. 4b). The “Line” and “Load” pressure sources used Fluigent LU-FEZ-2000 modules to control the pressures. The “Line” pressure source was set to 100 kPa. The “Load” pressure source was swept from 0 kPa to 50 kPa over the course of 300 s and the flow was recorded.

### Level shifter characterization

The pinout for the level shifter chip is given in Extended Data Figure 8j. Extended Data Figure 8k provides the setup used to demonstrate the level shifter (Fig. 2c). The “Supply” and “Input” pressure sources used a Fluigent LU-FEZ-7000 and a Fluigent LU-FEZ-2000 module respectively to control the pressure. The “Offset” pressure source was used to offset the pressure measurement and ensure an appropriate measurement range for the pressure sensor. The “Supply” pressure source was set to 250 kPa and the “Offset” pressure source was set to 150 kPa. The “Input” pressure source generated a sinusoidal waveform with an amplitude of 20 kPa, a baseline bias pressure of 80 kPa, and a period of 30 s. The output pressure waveform was recorded using a pressure sensor and plotted over 150 s (five periods).

The same setup (Extended Data Fig. 8k) was used to measure the level shifter shift amount and gain (Extended Data Fig. 4c-d). The “Supply” pressure source was set to 250 kPa and the “Offset” pressure source was set to 150 kPa. The “Input” pressure source was swept from 10 kPa to 90 kPa over the course of 240 s and the output pressure was recorded. The shift amount was determined by subtracting the output pressure from the pressure applied at the “Input” pressure source. The output pressure data was smoothed using a polynomial function of degree three to remove measurement noise, then the gain was calculated from the derivative. Note that this circuit operates in a common-drain configuration, and so the pressure gain is expected to be less than unity.

### NAND gate characterization

The pinout for the NAND gate is given in Extended Data Figure 9a. Extended Data Figure 9b provides the setup used to demonstrate the NAND gate (Fig. 2d). The “Supply” pressure source used a Fluigent LU-FEZ-7000 module to control the pressure. The “InHigh” and “InLow” pressure sources used two Fluigent LU-FEZ-2000 modules. The “Offset” pressure source used a Fluigent LU-FEZ-1000. “Switch1” and “Switch2” were Fluigent 2-switches (2SW002). The “Supply” pressure source was set to 150 kPa, the “Offset” pressure source was set to 100 kPa, the “InLow” pressure source was set to 125 kPa, and the “InHigh” pressure source was set to 175 kPa. Both “Switch1” and “Switch2” were set to toggle every 2.5 s, resulting in two square wave pressure signals with a period of 5 s. The switches were timed such that the two pressure waveforms had a 1.25 s phase delay between them. The output pressure signal was recorded over the course of 300 s.

The same setup (Extended Data Fig. 9b) was used to measure the NAND gate output dynamics (Extended Data Fig. 5a-b), revealing the maximum rate of change in the circuit output. The “Supply” pressure source was set to 150 kPa, the “InLow” pressure source was set to 125 kPa, and the “InHigh” pressure source was set to 175 kPa. “Switch1” was set to toggle every 2.5 s, while “Switch2” was maintained in the top position, connecting the “InB” port to the “InHigh” pressure source. The output pressure signal was recorded over the course of 300 s. Fifty-five individual rising and falling edges were overlaid and plotted.

Extended Data Figure 9c provides the setup used to measure the NAND gate transfer characteristics (Extended Data Fig. 5c-d). The “Supply” pressure source used a Fluigent LU-FEZ-7000 module to control the pressure. The “InputA” and “InputB” pressure sources used two Fluigent LU-FEZ-2000 modules. The “Offset” pressure source used a Fluigent LU-FEZ-1000. The “Supply” pressure source was set to 150 kPa and the “Offset” pressure source was set to 100 kPa. To measure the Input A transfer characteristics (Extended Data Fig. 5c), the “Input A” pressure source was swept from 125 kPa to 175 kPa over the course of 15 seconds while “Input B” was held high at 175 kPa. Subsequently, to measure the Input B transfer characteristics (Extended Data Fig. 5d), the “Input B” pressure source was swept from 175 kPa to 125 kPa over the course of 15 seconds while “Input A” was held high at 175 kPa. The output pressure signal was recorded as these sweeps were repeated ten times each. These transfer characteristics were overlaid and plotted.

### SR-latch characterization

The pinout for the SR-latch is given in Extended Data Figure 9d. Extended Data Figure 9e provides the setup used to demonstrate the SR-latch (Fig. 2e). The “Supply” pressure source used a Fluigent LU-FEZ-7000, the “InHigh” pressure source used a Fluigent LU-FEZ-2000, and the “Offset” pressure source used a Fluigent LU-FEZ-1000. “Switch1” and “Switch2” were Fluigent 2-switches (2SW002) normally in the open state. The “Supply” pressure source was set to 250 kPa, the “InHigh” pressure source was set to 165 kPa, and the “Offset” pressure source was set to 100 kPa. The latch was set by briefly closing and re-opening “Switch1” for the shortest period the Fluigent SDK would allow (0.5 s). The latch was then reset by briefly closing and re-opening “Switch2” for the shortest period the Fluigent SDK would allow. To demonstrate the memory of the latch (Fig. 2e), the output pressures were recorded as it was set and reset with arbitrarily varying time intervals between the set and reset operations.

The same setup (Extended Data Fig. 9e) was used to measure the SR-latch set and reset response (Extended Data Fig. 5e-f), revealing the response dynamics and speed of the circuit. The “Supply” pressure source was set to 250 kPa, the “InHigh” pressure source was set to 165 kPa, and the “Offset” pressure source was set to 100 kPa. The set and reset operations were done by briefly closing the switches as described above. In this fashion, the latch was alternatively set and reset every 2.5 s while the output pressures were measured over the course of 300 s. The resulting pressure signal consisted of sixty reset output edges (Extended Data Fig. 5e) and sixty set complementary edges (Extended Data Fig. 5f).

### Smart particle dispenser characterization

The concentration and ordering capabilities of the smart particle dispenser circuit were tested using a suspension of polystyrene microspheres in PBS. The suspension was prepared by adding 40 μm diameter polystyrene beads (Fluoro-Max Green 35-7B, Thermo-Fisher) to 50 mL of 1X PBS (Gibco PBS, Fisher Scientific) to achieve a final concentration of approximately 30 beads/mL.

The pinout for the particle trap is given in Extended Data Figure 9f. Extended Data Figure 9g provides the setup used to test the smart dispenser configured for particle concentration and ordering. The reservoir (green) connected to the “Part In” line of the trap was filled with the dilute polystyrene bead suspension and all other reservoirs were filled with PBS. The reservoirs connected to the “Supply” pressure source were 500 mL bottles, while all other reservoirs were P-CAP reservoirs from Fluigent. The “Supply” pressure source used a Fluigent LU-FEZ-7000 module to control the pressure. The “InHigh”, “OutLow”, and “Reference” pressure sources used Fluigent LU-FEZ-2000 modules to control the pressure. The “Sensor Offset” pressure source used a Fluigent LU-FEZ-1000 module to offset the pressure sensors, ensuring an appropriate measurement range. The tubing dimensions used for the resistances are provided in Table S1. The “Supply” pressure source was set to 250 kPa, the “InHigh” pressure source was set to 160 kPa, the “OutLow” pressure source was set to 140 kPa, the “Reference” pressure source was set to 150 kPa, and the “Sensor Offset” pressure source was set to 100 kPa.

All pressure sources remained constant during the entirety of the experiment, since all of the dynamic signal processing was performed by the microfluidic chip itself. Trapping events were consistently detected by a sharp rising edge in the *P_plug_* pressure signal, and additionally verified visually under a microscope. Between trapping events, the flow through the “Part In” line (*Q_in_*) was integrated to compute the input particle spacing volume, and the flow through the “Part Out” line (*Q_out_*) was integrated to compute the output particle spacing volume. The experiment was run for 230 trapping events before the “Supply” reservoirs of liquid to power the system were depleted.

## Supplementary information

### Text I: Calculation of intrinsic gain

While intrinsic gain was originally defined in the context of electronic transistors in terms of voltage and current^15^, we may follow an analogous derivation to define the intrinsic gain for a microfluidic transistor in terms of pressure and flow. For a microfluidic transistor where the flow *Q* is a function of the pressures *P_SD_* and *P_GS_* applied across its terminals, the transconductance *g_m_* is given by:

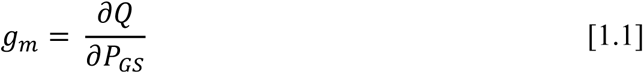

And the output impedance *r*_0_ is given by:

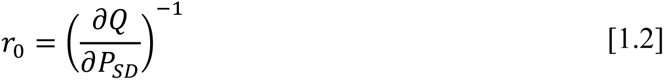

Then the dimensionless intrinsic gain *A*_0_ is given by:

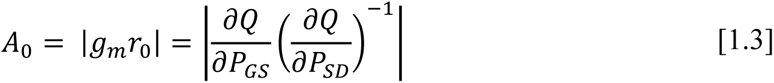

Note that this is analogous to the formula used in electronics for field-effect transistors, substituting pressure and flow for voltage and current^15^.

### Text II: Shapiro number for rectangular channels

In his seminal work describing flow-limitation, Ascher Shapiro mathematically modeled the flow of an internal incompressible Newtonian fluid through a thin-walled deformable tube^17^. For this system, Shapiro defined a “characteristic wave propagation speed” *c* by the following:

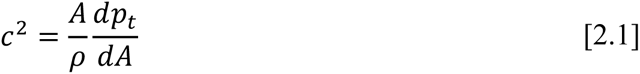

Where *A* is a characteristic cross-sectional area of the tube, and *ρ* is the fluid density. The term 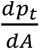 couples structural deformation of the tube to the fluid flow. In previous studies, this term has been deduced based on the “tube law” for the system, which is the relationship between the cross-sectional area of the tube and the transmural pressure *p_t_* across its walls. Typically, if the internal pressure of the tube is held constant, increasing the external pressure will cause the tube to deform and cause its cross-sectional area to drop.

While the empirically derived tube law relationship was originally used to describe the deformation of thin-walled cylindrical tubes, here we consider the deformation of a square piece of thin membrane over a channel with a rectangular cross-section (Fig. 1a). The reciprocal hydraulic compliance of this membrane-channel fluidic system can be derived by plate theory as^12^:

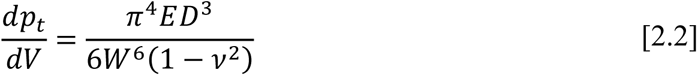

Where *V* is the volume of fluid in the channel under the membrane, *W* is the characteristic length scale of the square membrane, *D* is the membrane thickness, *E* is the Young’s modulus of the membrane material, and *v* is the Poisson ratio of the membrane material. Dividing both sides by the length of the square membrane, we obtain the following characteristic “tube law” for a channel with a deformable square membrane:

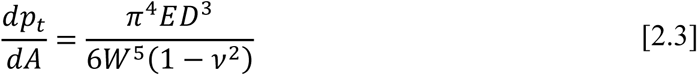

Substituting this into Equation 2.1 we obtain the following expression for the characteristic wave speed *c*:

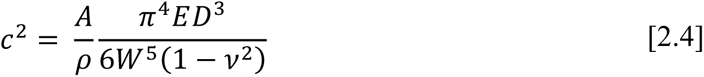

The Shapiro number *S* for this system is then simply the ratio of the characteristic fluid velocity to the characteristic wave speed of the channel. In terms of the flow rate *Q*, this is given by:

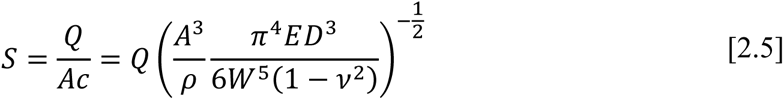

For the microfluidic transistor characterized in Fig. 1c, the channel width *W* is 500 μm, the characteristic cross-sectional area *A* is 0.0275 mm^2^, the membrane thickness *D* is 20 μm, the membrane Poisson’s ratio *v* is 0.5, the Young’s modulus *E* is 550 kPa, and the fluid density *ρ* is 1.01 g/mL^28,29^. We may then use the characteristic curve measurements to compute the Shapiro number directly from the measured flow rate (Extended Data Fig. 2). Note that in this analysis we only consider the curve where *P_GS_* = 0, which is the case analyzed by Shapiro.

The Shapiro number delineates a critical transition in the behavior of the membrane-channel system (Extended Data Fig. 2). When the Shapiro number is much less than one, the deformation of the membrane does not significantly restrict flow, and the channel exhibits flow-pressure relationships as predicted by the Poiseuille equation. When the Shapiro number is greater than one, the deformation of the membrane significantly restricts flow, and the phenomenon of flow-limitation takes place^23^.

## Acknowledgements

We are very grateful to Octavio Hurtado for his insights and assistance with microfabrication. The project described was supported by award Number T32GM007753 from the National Institute of General Medical Sciences. The content is solely the responsibility of the authors and does not necessarily represent the official views of the National Institute of General Medical Sciences or the National Institutes of Health.

## Declarations

### Author contributions (CRediT Model)

Conceptualization: KAG, MT

Methodology: KAG, MT

Investigation: KAG, AM, BRM, JFE

Visualization: KAG

Funding acquisition: KAG, AM, JFE, MT

Project administration: MT

Supervision: JFE, MT

Writing – original draft: KAG, AM

Writing – review & editing: KAG, AM, JFE, MT

### Availability of data and materials

All relevant data are included in the article and/or its supplementary information files. No specialized code or algorithms were used to analyze the data in the current study.

### Additional Information

Supplementary Information is available for this paper. Correspondence and requests for materials should be addressed to Mehmet Toner.

**Extended Data Fig. 1.**
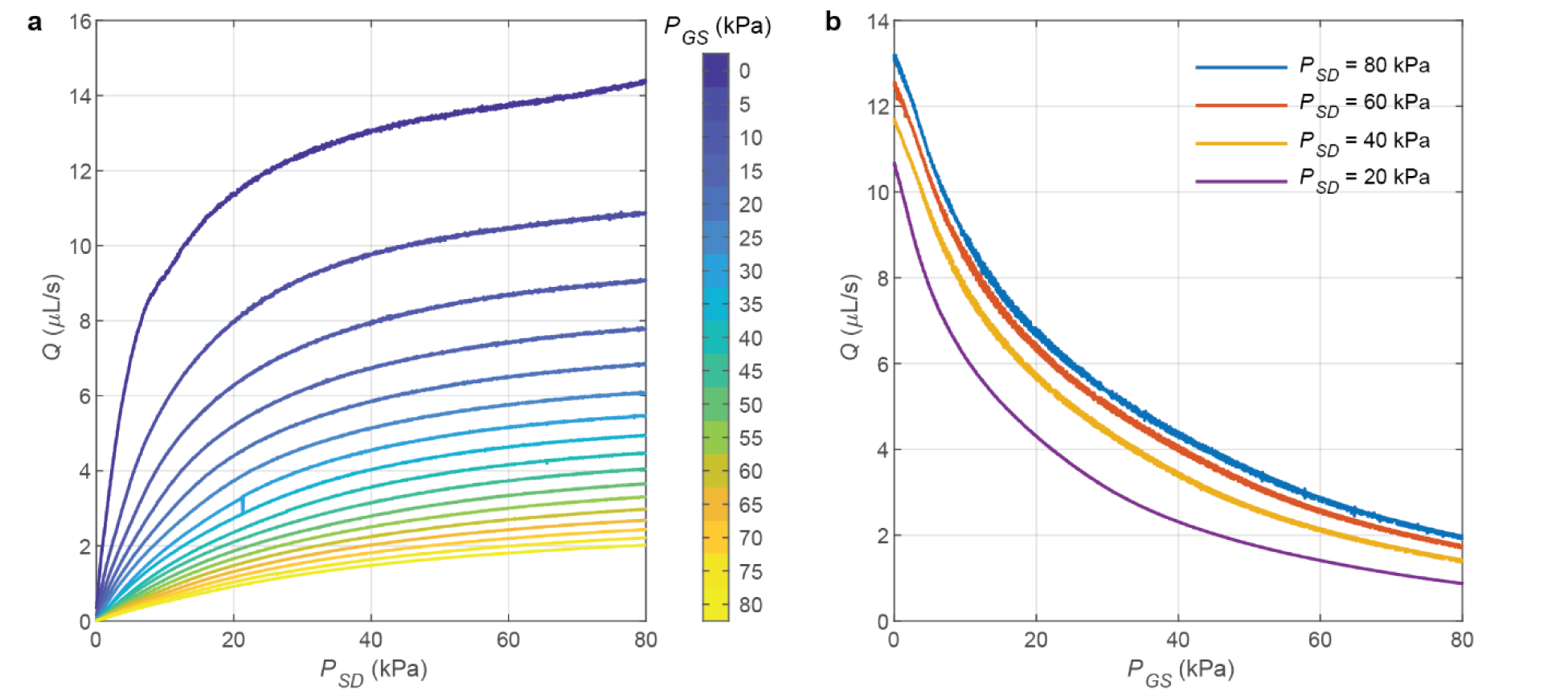
Additional characterization of a microfluidic transistor. **a,** Experimentally measured characteristic curves for a single transistor. At large values of *P_SD_*, the device exhibits a saturation-like phenomenon due to flow-limitation. The flow rate can be modulated by applying a pressure *P_GS_*. **b,** Transconductance characteristics for a microfluidic transistor. The high slopes of the curves at low *P_GS_* indicate that the device has a high transconductance suitable for analog amplification.

**Extended Data Fig. 2.**
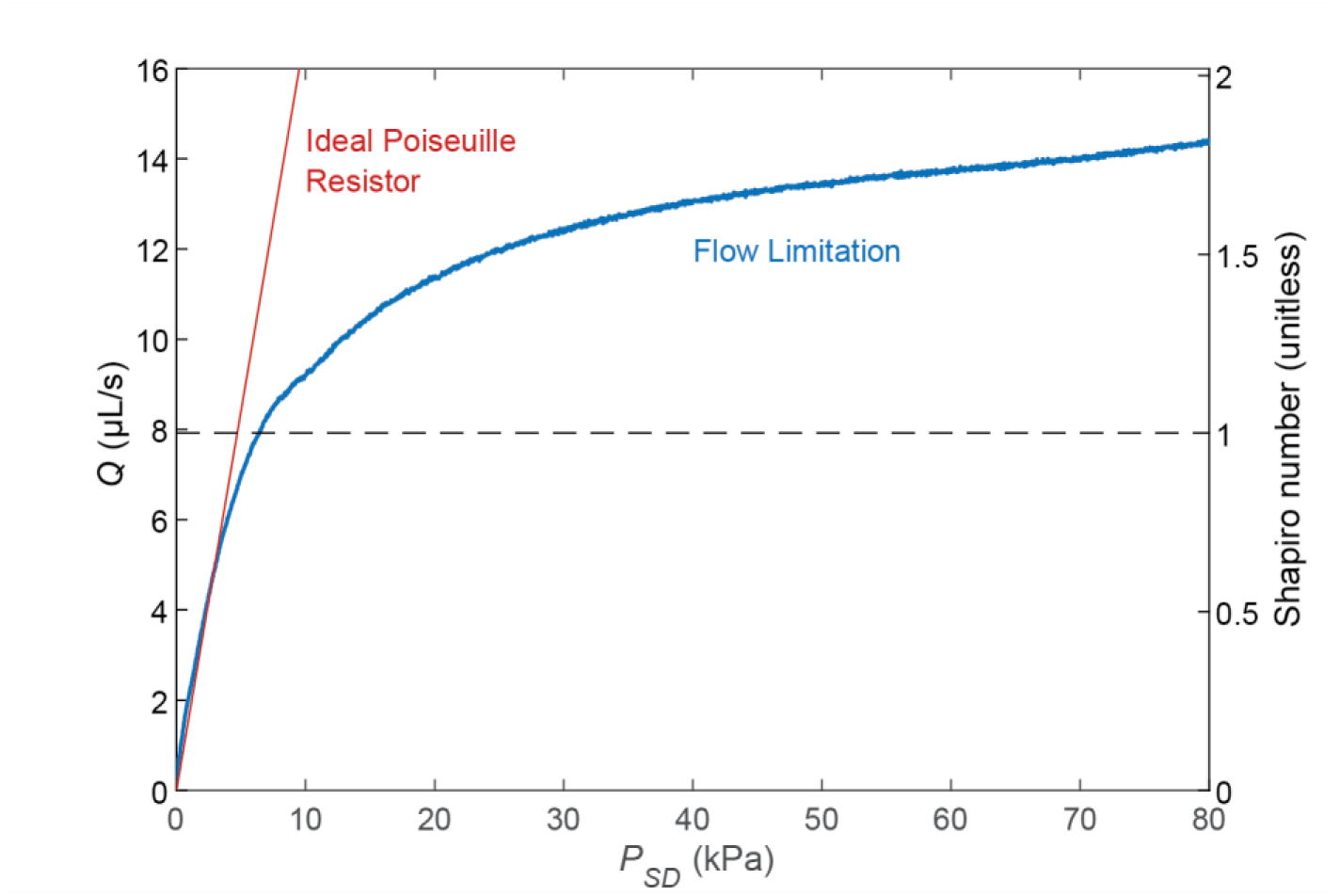
Microfluidic transistors exhibit flow limitation as the Shapiro number exceeds one. In the regime where the Shapiro number is less than one, the flow-pressure characteristics of the microfluidic transistor (blue) follow the linear relationship predicted by the Poiseuille equation (red). As the Shapiro number exceeds one (dashed line), the flow-pressure characteristics deviate, and the system exhibits flow limitation.

**Extended Data Fig. 3.**
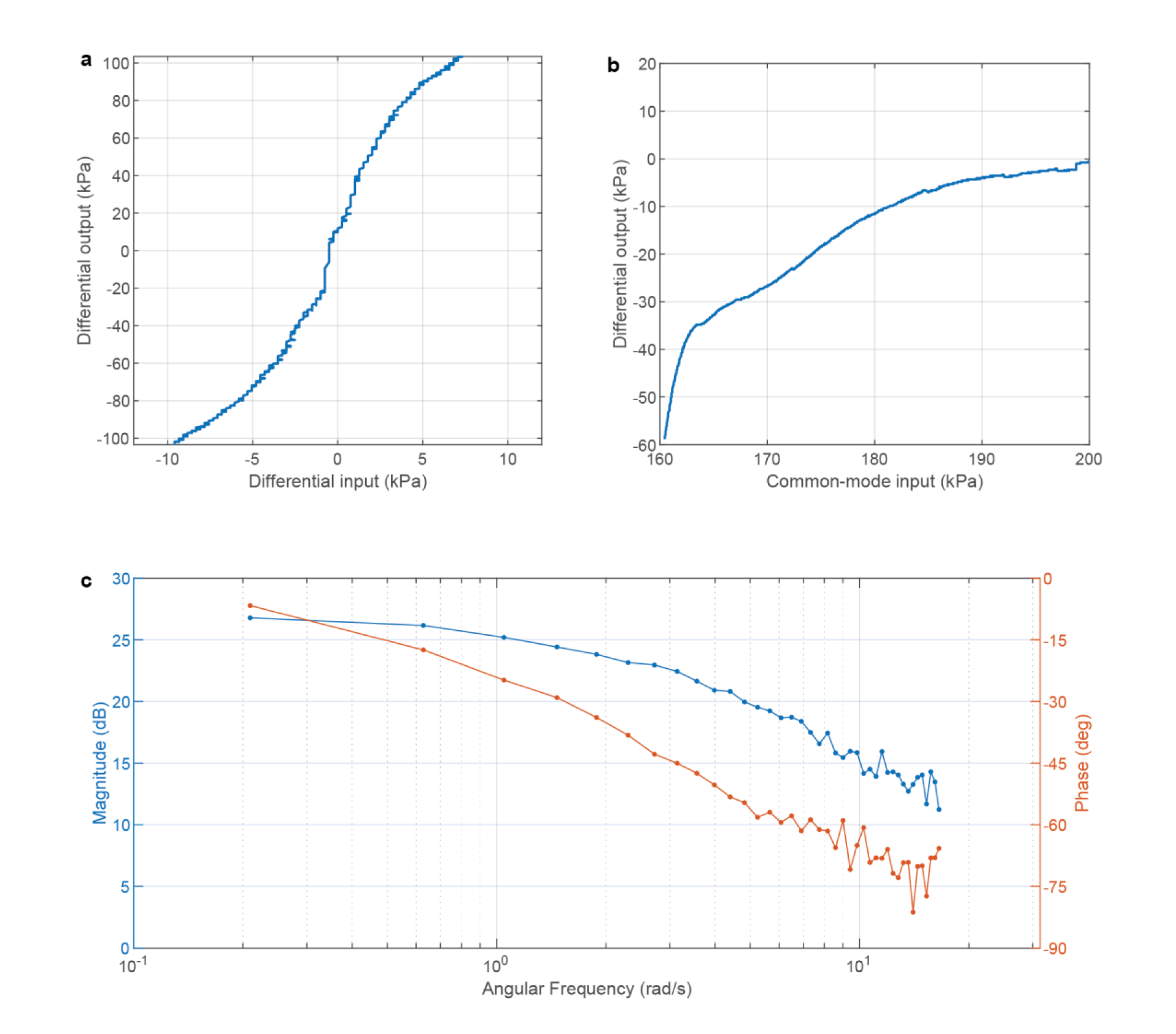
Additional characterization of the differential amplifier. **a,** Distortion (Transfer) characteristics of the differential amplifier. The differential amplifier provides relatively linear amplification for amplitudes below 5 kPa. As in electronics, application of negative feedback can further linearize this output^16^. **b,** Common-mode rejection of the differential amplifier. The differential output is less sensitive to common-mode changes when biased around 190 kPa. **c,** Frequency response of the differential amplifier. Fourier analysis of the first 40 odd-harmonics of a square wave input show a low-frequency gain of approximately 27 dB.

**Extended Data Fig. 4.**
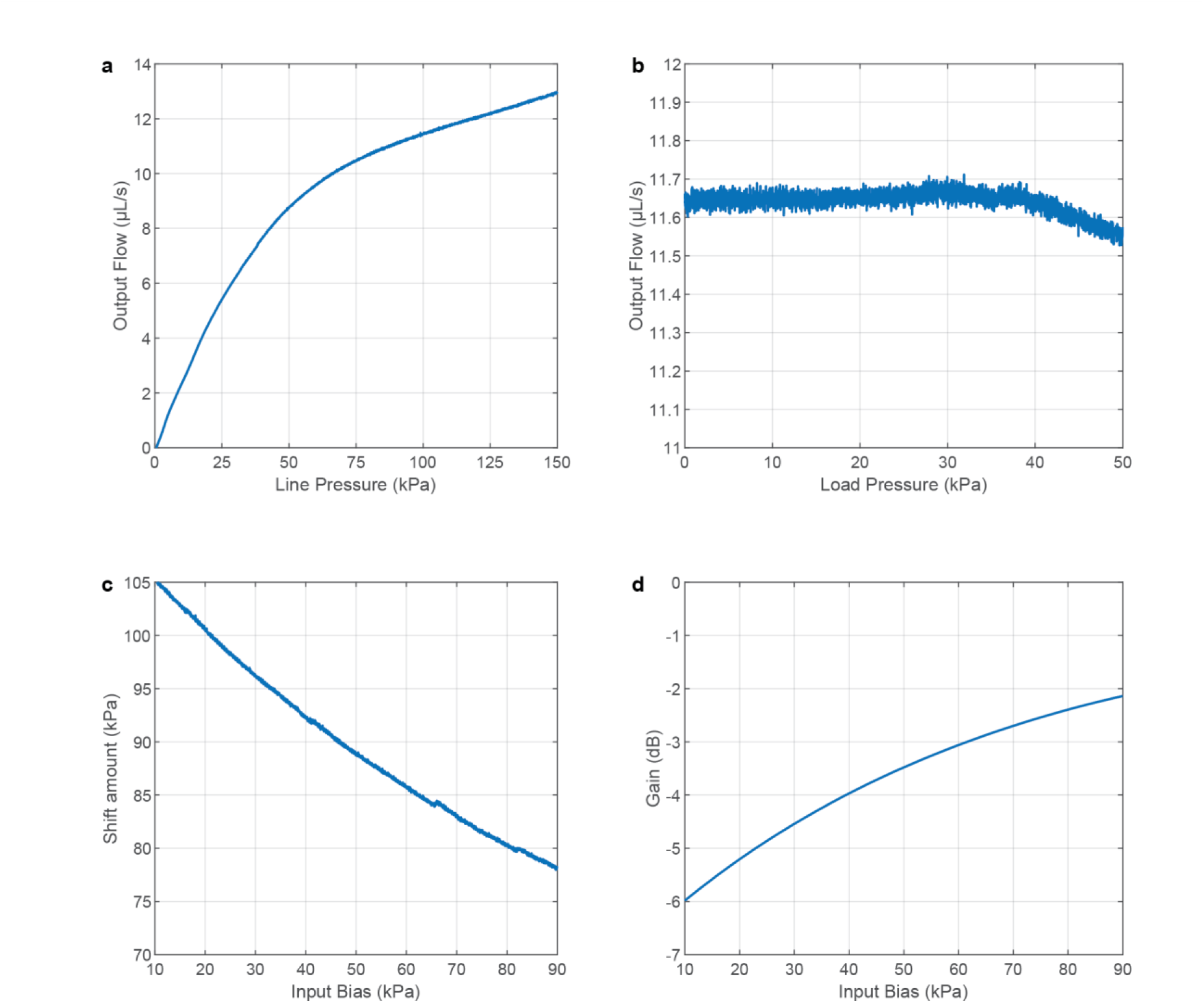
Additional characterization of the flow regulator and the level shifter. **a,** Line regulation of the flow regulator through a 2 kPa s μL^-1^ load. The output flow is less sensitive to changes to the line pressure above 75 kPa. **b,** Load regulation of the flow regulator. With a line pressure of 100 kPa, the output flow is insensitive to changes in the load up to 50 kPa. **c,** Pressure shift of the level shifter. The level shifter is capable of shifting signals by over 80 kPa. **d,** Gain of the level shifter. Depending on the bias, there is a small drop in the amplitude of output signal. Note that since this circuit operates in a common-drain configuration, the gain in decibels is expected to be negative.

**Extended Data Fig. 5.**
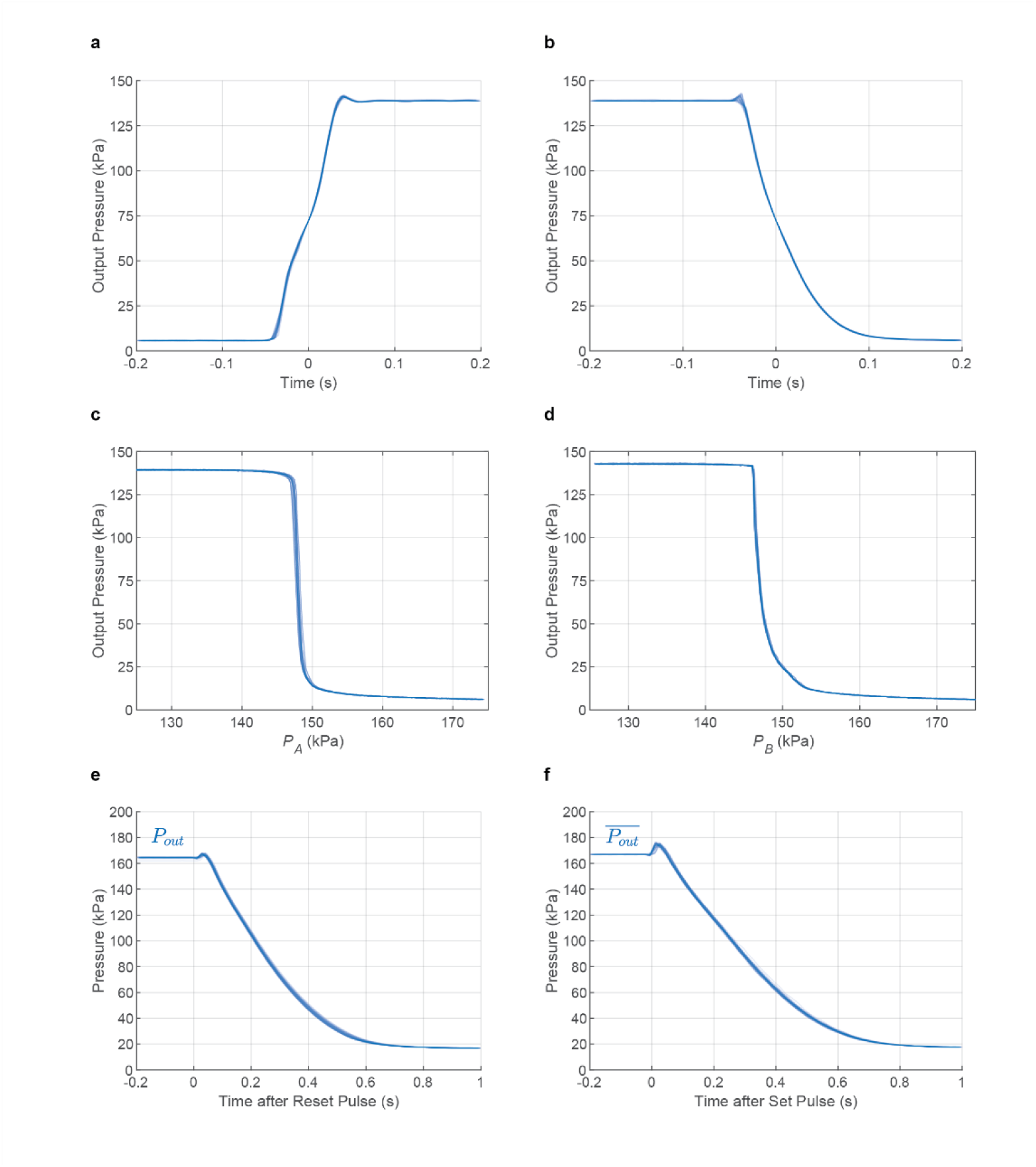
Additional characterization of the NAND gate and the SR Latch. **a-b,** Output transition and dynamics of the NAND gate. Input signal B was set to high while input signal A was toggled between high and low, causing the output to toggle between low and high. The transition for n=55 individual rising (**a**) and falling (**b**) output edges are overlaid, showing rise and fall times of less than 100 ms with good repeatability. **c-d,** Transfer characteristics of the NAND gate. **c,** Input A was swept across a range of pressures while input B was held high, and the output signals for n=10 sweeps were overlaid. **d,** Input B was then swept while input A was held high, and the output signals for n=10 sweeps were overlaid. The sharp transitions between logic levels indicate a large noise margin for digital signals. **e,** Reset response dynamics of the SR-latch. After initializing the latch to a high output state (*P_out_* high and 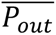 low), a reset pulse flipped the output *P_out_* low within 800 ms. The overlaid output for n=60 such events demonstrates repeatability. **f,** Set response dynamics of the SR-latch. After initializing the latch to a low output state (*P_out_* low and 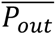 high), a set pulse flipped the complementary output 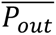 low within 800 ms. The overlaid output for n=60 such events demonstrates repeatability.

**Extended Data Fig. 6.**
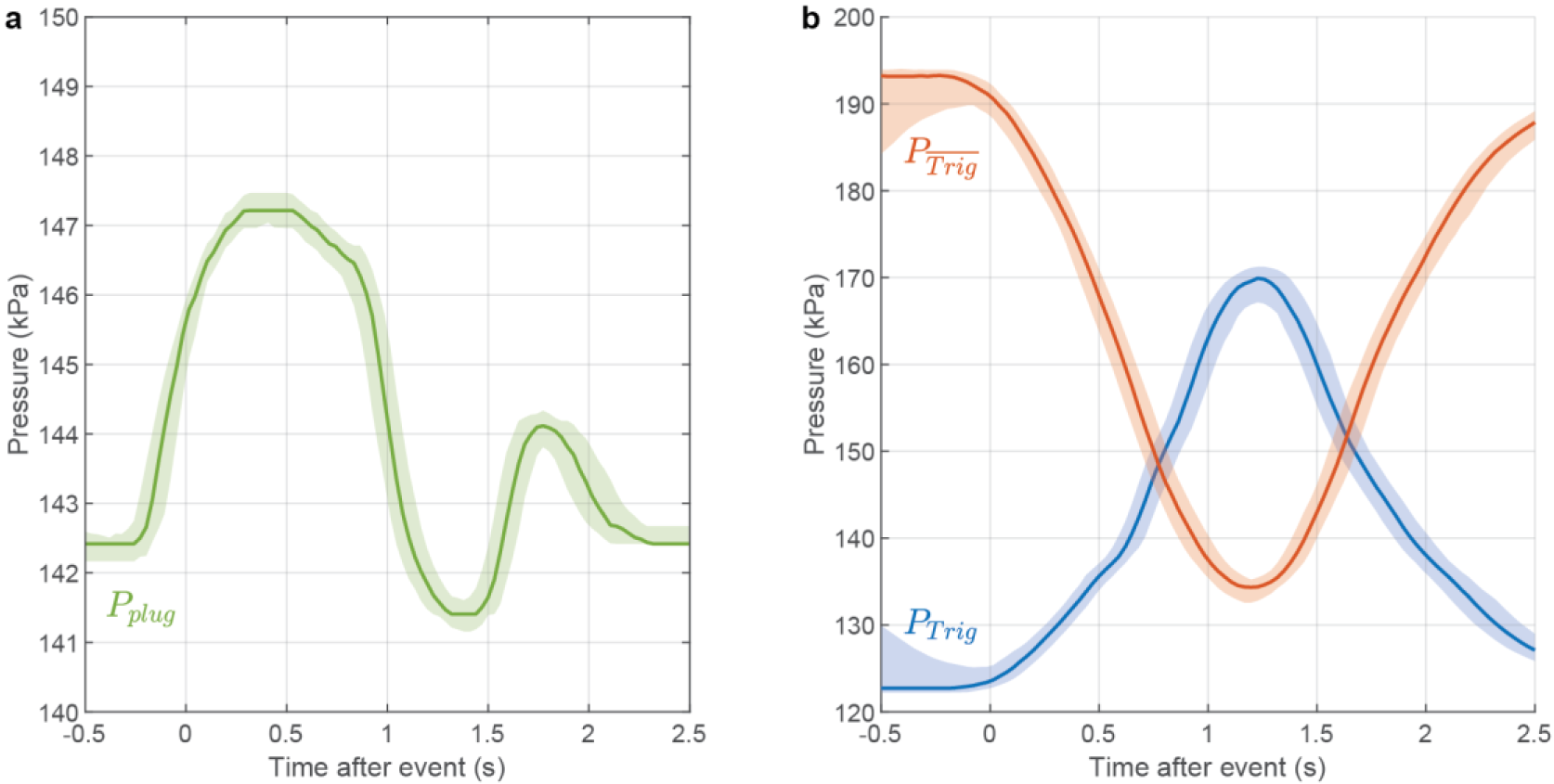
Repeatability of the smart particle dispenser. **a,** Collated plug signals for individual events. After a particle is trapped, the pressure upstream of the trap (*P_plug_*) rises in a repeatable fashion. The *P_plug_* pressure signal for n=230 trapping events were aggregated, and a pointwise median signal with 90% and 10% quantile bands were plotted. **b,** Collated trigger signals for individual events. The amplifier and the level shifters process the *P_plug_* signal to produce the trigger signals *P_Trig_* and 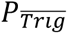. These signals for n=230 trapping events were aggregated, and a pointwise median signal with 90% and 10% quantile bands are plotted.

**Extended Data Fig. 7.**
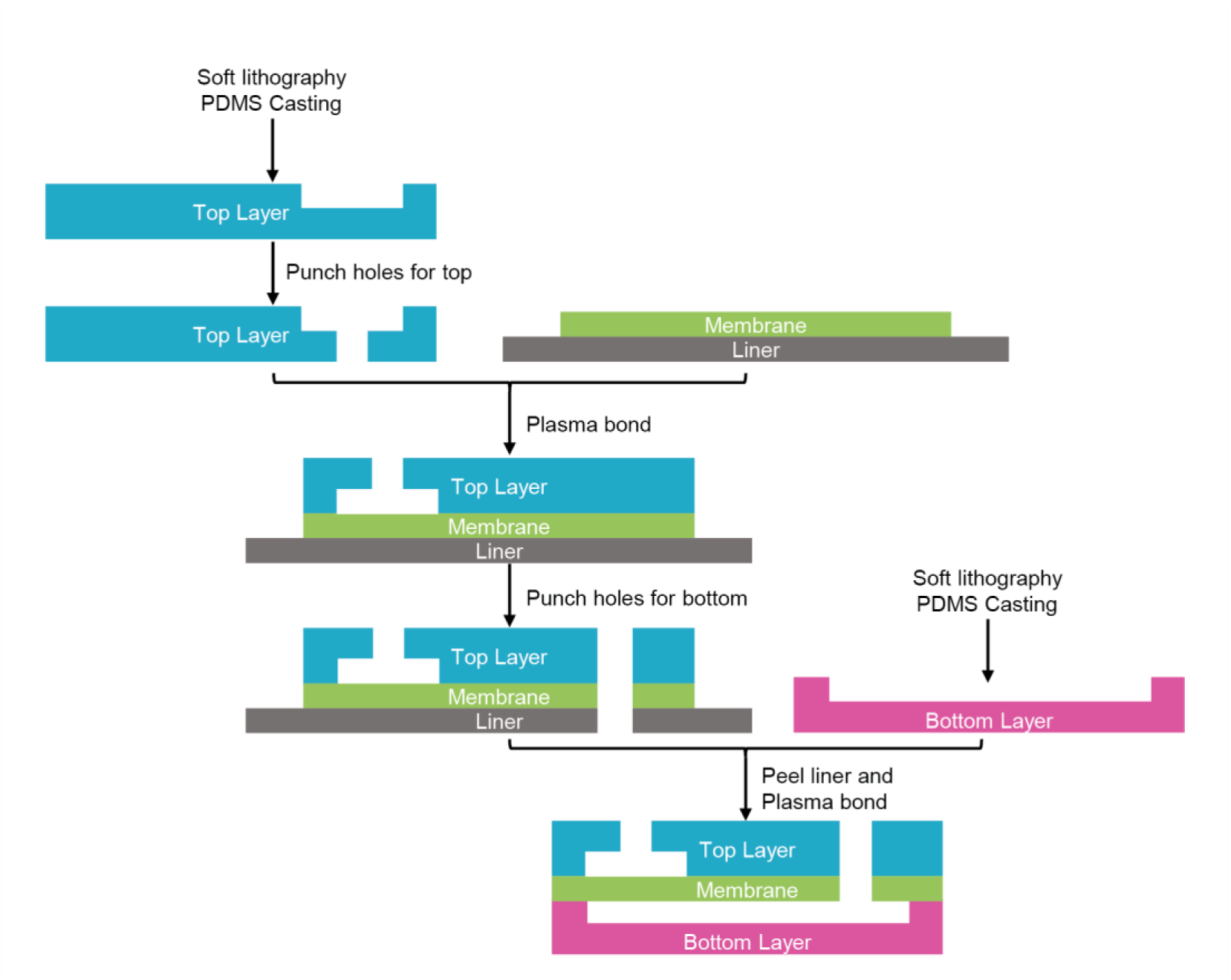
Assembly of multilayer microfluidic transistor-based circuits. Soft-lithography techniques are used to fabricate the top and bottom PDMS layers. Top layer ports are punched into the top PDMS layer. Then, the layer is bonded with a thin silicone membrane under oxygen plasma. Ports for the bottom layer are then punched into the top layer-membrane assembly. The assembly is aligned by hand and finally bonded with the bottom layer under oxygen plasma.

**Extended Data Fig. 8.**
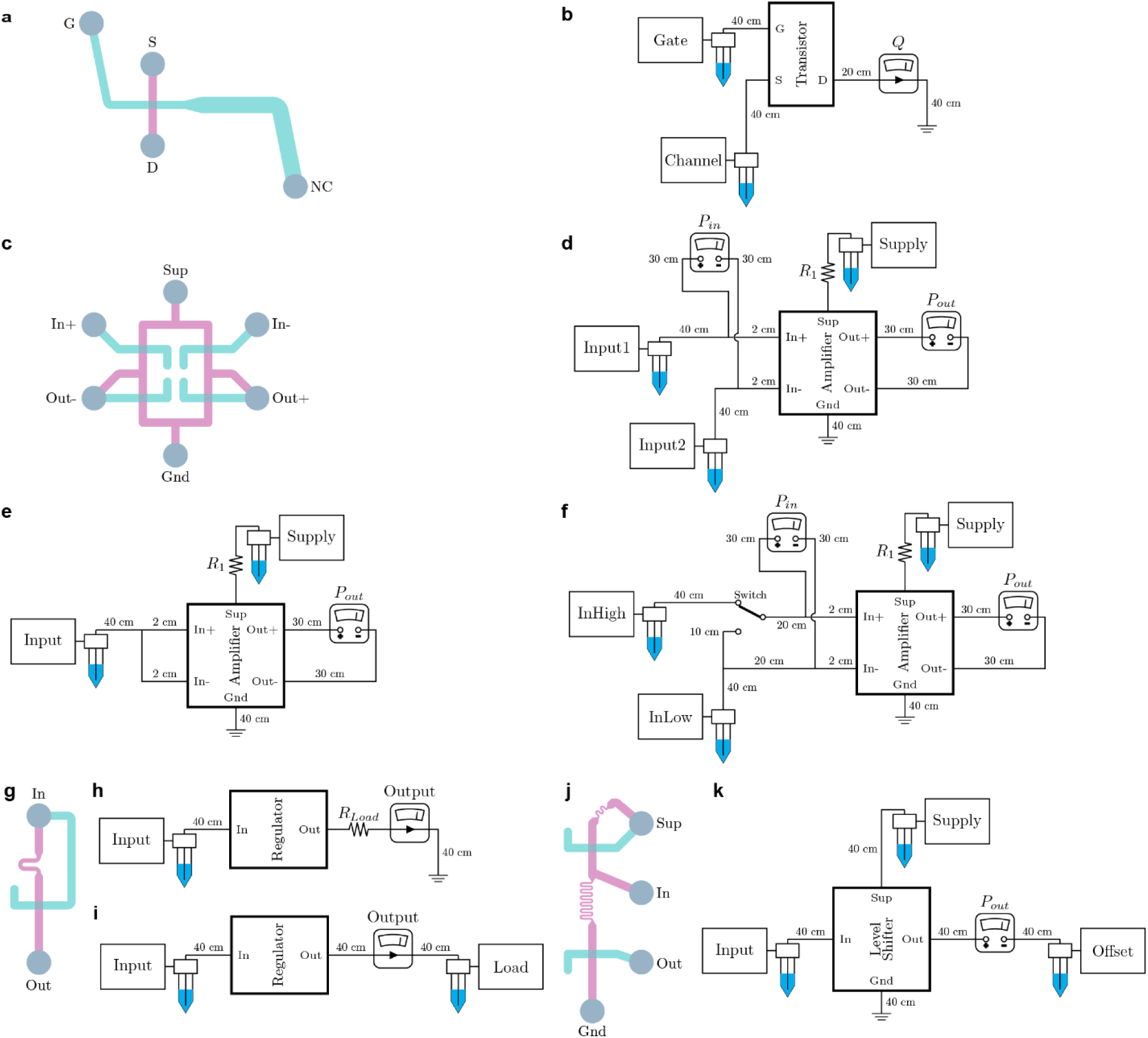
Setups for single transistor, amplifier, flow regulator, and level shifter measurements. Relevant component details such as geometry and resistance values are provided in Extended Data Table 1. **a,** Pinout diagram of single transistor chip (NC: no connection) with channel layers colored magenta and teal. **b,** Fluidic setup for characteristic curves and transconductance curve measurements. **c,** Pinout diagram of amplifier chip with channel layers colored magenta and teal. **d,** Fluidic setup for demonstration and distortion measurements. **e,** Fluidic setup for common-mode rejection measurements. **f,** Fluidic setup for Frequency response (Bode plot) measurements. **g,** Pinout diagram of flow regulator chip with channel layers colored magenta and teal. **h,** Fluidic setup for demonstration and line regulation measurements. **i,** Fluidic setup for load regulation measurements. **j,** Pinout diagram of level shifter chip with channel layers colored magenta and teal. **k,** Fluidic setup for demonstration, shift amount, and gain measurements.

**Extended Data Fig. 9.**
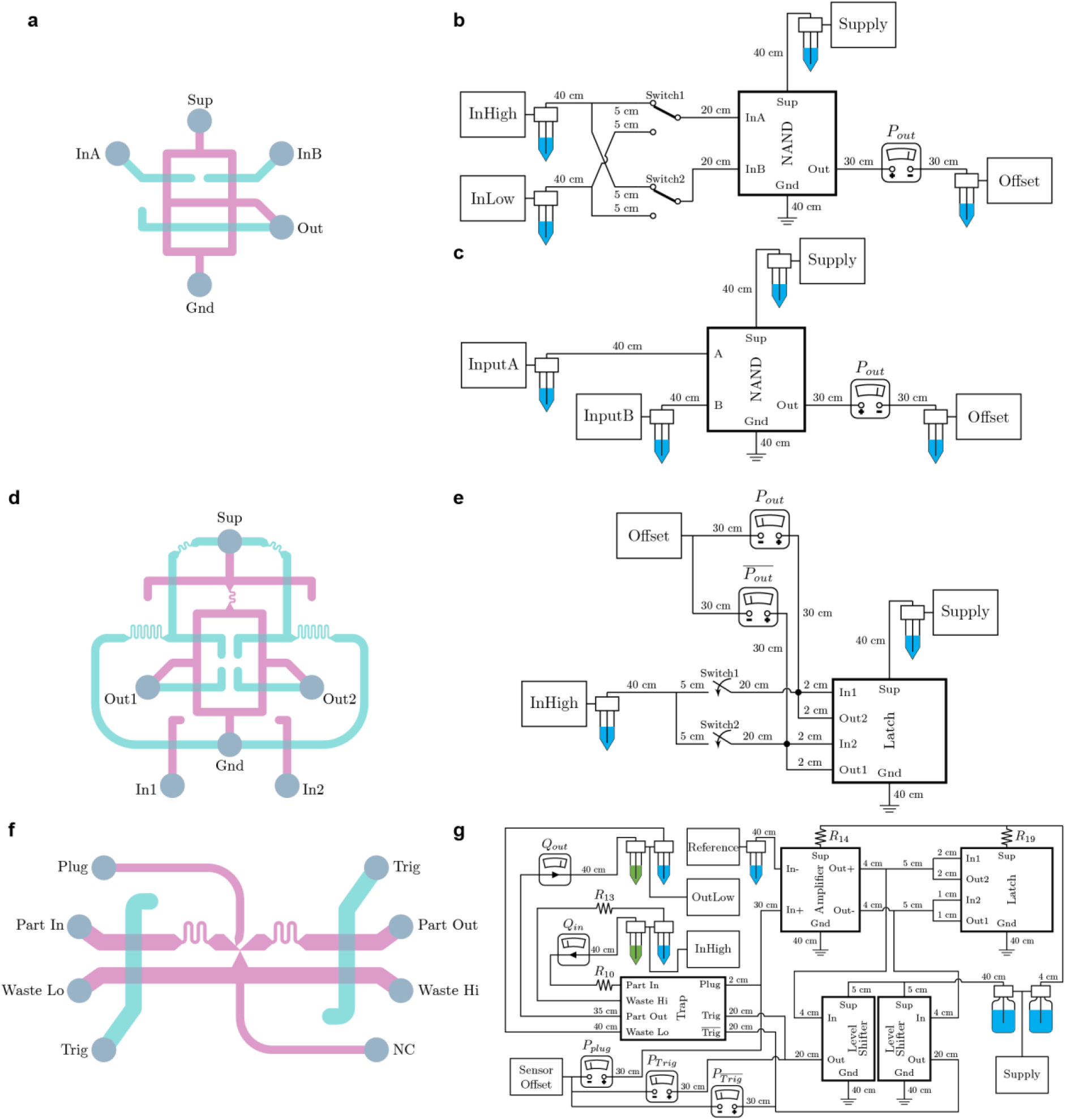
Setups for NAND gate, SR-latch, and smart particle dispenser measurements. Relevant component details such as geometry and resistance values are provided in Extended Data Table 1. **a,** Pinout diagram of NAND gate chip with channel layers colored magenta and teal. **b,** Fluidic setup for demonstration and output dynamics measurements. **c,** Fluidic setup for transfer characteristics measurements. **d,** Pinout diagram of SR-latch chip with channel layers colored magenta and teal. **e,** Fluidic setup for demonstration and response dynamics measurements. **f,** Pinout diagram of particle trap chip (NC: no connection) with channel layers colored magenta and teal. **g,** Fluidic setup for particle dispenser ordering and concentration. Several blocks are used for signal processing. The “Supply” sources used 500mL bottles as reservoirs. Green reservoirs hold bead suspensions. In this configuration the Trig and Sense signals are directly connected to each other to perform particle ordering and concentration.

**Extended Data Table 1.**
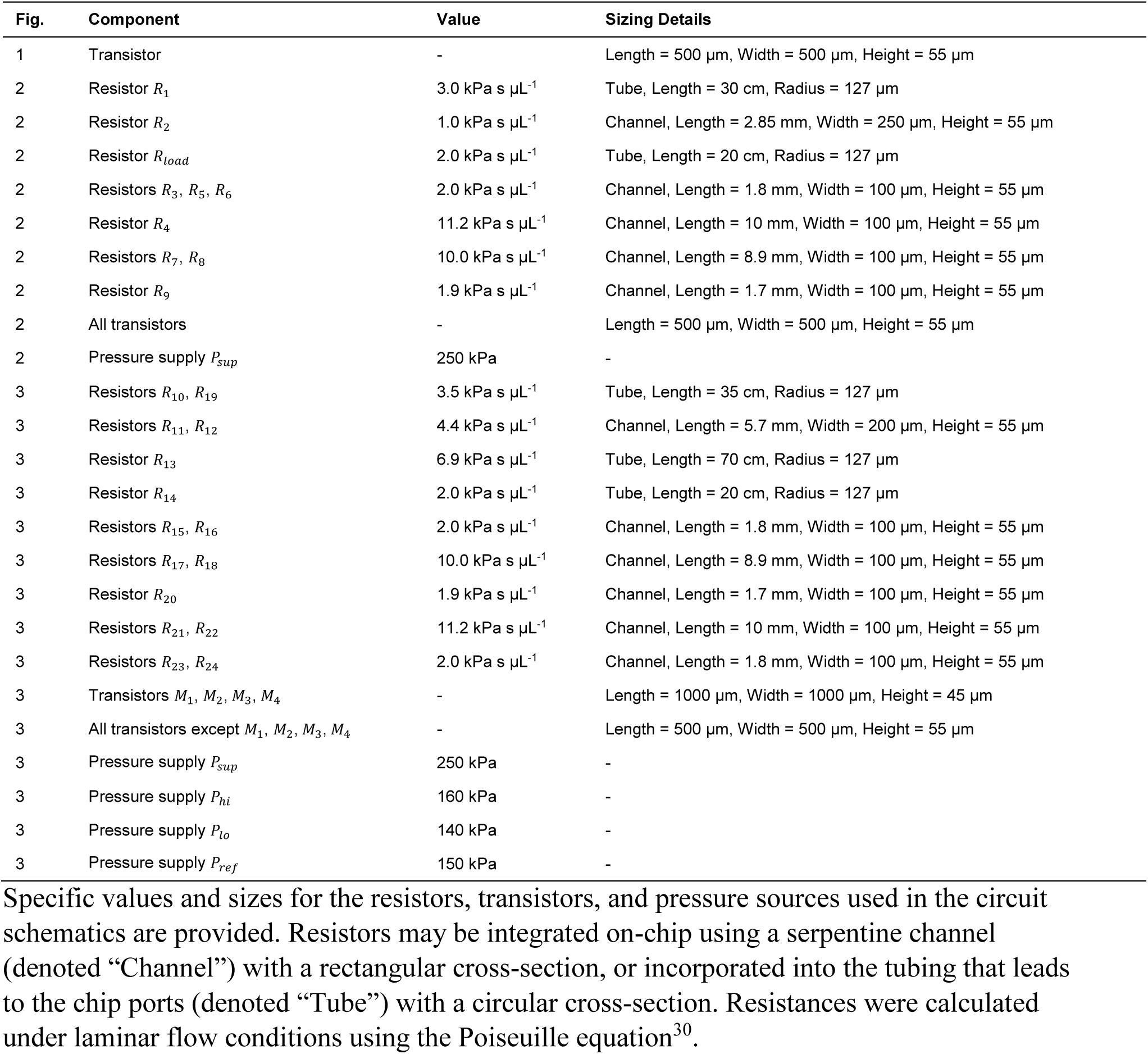
Circuit schematic component details.

